# A genetic screen using the *Drosophila melanogaster* TRiP RNAi collection to identify metabolic enzymes required for eye development

**DOI:** 10.1101/582940

**Authors:** Rose C. Pletcher, Sara L. Hardman, Sydney F. Intagliata, Rachael L. Lawson, Aumunique Page, Jason M. Tennessen

## Abstract

The metabolic enzymes that compose glycolysis, the citric acid cycle, and other pathways within central carbon metabolism have emerged as key regulators of animal development. These enzymes not only generate the energy and biosynthetic precursors required to support cell proliferation and differentiation, but also moonlight as regulators of transcription, translation, and signal transduction. Many of the genes associated with animal metabolism, however, have never been analyzed in a developmental context, thus highlighting how little is known about the intersection of metabolism and development. Here we address this deficiency by using the *Drosophila* TRiP RNAi collection to disrupt the expression of over 1,100 metabolism-associated genes within cells of the eye imaginal disc. Our screen not only confirmed previous observations that oxidative phosphorylation serves a critical role in the developing eye, but also implicated a host of other metabolic enzymes in the growth and differentiation of this organ. Notably, our analysis revealed a requirement for glutamine and glutamate metabolic processes in eye development, thereby revealing a role of these amino acids in promoting *Drosophila* tissue growth. Overall, our analysis highlights how the *Drosophila* eye can serve as a powerful tool for dissecting the relationship between development and metabolism.

## INTRODUCTION

The fruit fly *Drosophila melanogaster* has emerged as a powerful model for investigating the metabolic mechanisms that support animal growth and development. In this regard, a key advantage of studying metabolism in the fly is that the disruption of an individual metabolic reaction often induces a specific phenotype, thus revealing energetic and biosynthetic bottlenecks that influence cell growth, proliferation, and differentiation. For example, mutations that disrupt activity of the citric acid cycle (TCA cycle) enzymes Isocitrate Dehydrogenase 3b (Idh3b) and Malate Dehydrogenase 2 (Mdh2) prevent the larval salivary glands from dying at the onset of metamorphosis (Wang *et al*. 2008; Wang *et al*. 2010; Duncan *et al*. 2017). These observations suggest that the salivary glands are uniquely dependent on the TCA cycle to activate the cell death program and reveal an unexpected relationship between central carbon metabolism and metamorphosis. Such phenotype-driven studies are essential for investigating how metabolism and development are coordinated during the fly life cycle.

The *Drosophila* eye has long served as a powerful model for both metabolism and development (for reviews, see Dickinson and Sullivan 1975; Kumar 2018). Many of the earliest genetic studies conducted in the fly were based upon genes such as *vermillion*, *cinnabar*, and *rosy*, which control eye pigmentation and encode enzymes involved in tryptophan and purine metabolism (Lindsley and Zimm 1992). Similarly, classic work by Beadle and Ephrusi used transplantation experiments to demonstrate that ommochromes are synthesized in larval peripheral tissues and transported into the eye (Beadle and Ephrussi 1936), thus revealing that metabolism is systemically coordinated during development. Nearly a century later, the *Drosophila* eye still serves as an essential tool for studying developmental metabolism – a fact that is best illustrated by a finding from Utpal Banerjee’s lab. In a classic demonstration of how unbiased screens can identify unexpected developmental regulators, members of the Banerjee lab discovered that the *Drosophila* gene *CoVa* (FBgn0019624; also known as *tenured* and *COX5A*), a subunit of Complex IV within the electron transport chain (ETC) is essential for normal eye development (Mandal *et al*. 2005). While such a discovery could have been easily discounted as the disruption of a housekeeping gene, characterization of *CoVa* mutants demonstrated that reduced oxidative phosphorylation (OXPHOS) induces a G1 cell-cycle arrest during the second mitotic wave (Mandal *et al*. 2005). Moreover, this phenomenon was found to be orchestrated by the metabolic sensor AMPK, which responds to the decreased ATP levels present within *CoVa* mutant cells by activating p53 and lowering Cyclin E levels (Mandal *et al*. 2005). These studies of *CoVa* function, together with similar studies of other electron transport chain (ETC) subunits and mitochondrial tRNAs (Mandal *et al*. 2005; Liao *et al*. 2006), demonstrate that the eye can be used to efficiently understand how metabolism is integrated with developmental signaling pathways.

Here we use the *Drosophila* TRiP RNAi collection to identify additional metabolism-associated genes that influence eye development. Our screen used the *eyes absent composite enhancer-GAL4* (*eya composite-*GAL4) driver to induce expression of 1575 TRiP RNAi transgenes (representing 1129 genes) during development of the eye imaginal disc (Weasner *et al*. 2016). This analysis not only confirmed previous findings that genes involved in OXPHOS are essential for eye development, but also uncovered a role for glutamate and glutamine metabolism within this tissue. Moreover, we identified several poorly characterized enzymes that are essential for normal eye formation, thus hinting at novel links between metabolism and tissue development. Overall, our genetic screen provides a snapshot of the biosynthetic and energetic demands that the development of a specific organ imposes upon intermediary metabolism.

## METHODS

### Drosophila Strains and Husbandry

Fly stocks and crosses were maintained at 25° C on Bloomington *Drosophila* Stock Center (BDSC) food. All genetic crosses described herein used *eya composite-GAL4* to induce transgene expression (a kind gift from Justin Kumar’s lab, Weasner *et al*. 2016). The TRiP RNAi lines used in this study were selected by searching the BDSC stock collection using a previously described list of metabolism-associated genes (Tennessen *et al*. 2014; Perkins *et al*. 2015). All strains used in this study are available through the BDSC.

### Genetics Crosses and Phenotypic Characterization

Five adult male flies from each TRiP stock was crossed with five *w^1118^; eya composite-GAL4* adult virgin females. For each cross, F1 progeny heterozygous for both *eya composite-GAL4* and the *UAS-RNAi* transgene were scored for eye phenotypes within three days of eclosion. Eyes were scored for the following phenotypes: rough, glossy, small, no eye, misshaped, overgrowth, necrosis, abnormal pigmentation, and lethality prior to eclosion. Whenever possible, at least 20 adults were scored from during this screen. Any TRiP stock that produced a phenotype during the initial analysis was reanalyzed using the same mating scheme described above and twenty flies of each sex were scored. In some instances, expression of the TRiP transgene induced a lethal or semi-lethal phenotype prior to eclosion, thus limiting the number of animals that could be scored in our analysis. To avoid confirmation bias, each cross was only labeled with the BDSC strain number and the genotype was revealed only after phenotypic characterization.

### Statistical Analysis of Pathway Enrichment

Genes were assigned to individual pathways based on the metabolic pathways annotated within the Kyoto Encyclopedia of Genes and Genomes (KEGG) pathways (Kanehisa 2017; Kanehisa *et al*. 2019). Enrichment analyses for individual metabolic pathways were conducted using Fisher’s exact test with the P value being calculated using two tails. For the purpose of these calculations, expression of 165 transgenes induced a phenotype while the expression of 1410 transgenes had no effect on eye development.

## DATA AVAILABILITY

All TRiP lines used in this study are available from the BDSC. The *eya composite-GAL4* strain is available upon request. Data generated in this study have been uploaded to the RNAi Stock Validation and Phenotype (RSVP) database, which is publically available through the DRSC/TRiP Functional Genomics Resources website. The full list of TRiP stocks used in our analysis can be found in Supplemental Table 2 and the strains analyzed in the secondary screen are listed in Supplemental Table 3. To facilitate replication of our study, all supplemental tables contain both the BDSC and FlyBase identification numbers (Fbgn).

## RESULTS

To identify metabolic processes involved in eye development, we used the *eya composite*-*GAL4* driver to express TRiP RNAi transgenes that target metabolism-associated genes (Supplemental Table 1). Since this GAL4 driver promotes transgene expression in the eye imaginal disc from the L2 stage until after the morphogenetic furrow moves across the eye field (Weasner *et al*. 2016), our screen of 1575 TRiP RNAi lines was designed to identify metabolic processes required for the proliferation and differentiation of cells within this organ. Of the RNAi transgenes examined, 198 induced an eye phenotype in the initial screen and 165 subsequently generated reproducible phenotypes (Supplemental Tables 2 and 3).

Among the TRiP lines that consistently induced an eye phenotype were a number of positive controls. Notably, the RNAi transgene that targets *CoVa* (BDSC 27548) induced glossy-eye phenotype (Figure 1A-B; Supplemental Tables 2 and 3), thus phenocopying the eye defect associated with *CoVa* mutant clones (Mandal *et al*. 2005). Moreover, TRiP RNAi lines that interfere with expression of *Insulin Receptor* (BDSC 35251; FBgn0283499, REF), *PI3K* (BDSC 27690; FBgn0015279), *Akt* (BDSC 31701 and 33615; FBgn0010379), *Tor* (BDSC 34639; FBgn0021796), and *raptor* (BDSC 31528, 31529, and 34814; FBgn0029840) resulted in either small or misshapen eyes (Figure 1C; Supplemental Tables 2 and 3). Similarly, RNAi-induced depletion of the negative growth regulators *PTEN* (BDSC 25967 and 33643; FBgn0026379), *Tsc1* (BDSC 52931 and 54034; FBgn0026317) and *Tsc2* (BDSC 34737; FBgn0005198) induced an overgrowth phenotype (Figure 1D; Supplemental Tables 2 and 3). Our findings are consistent with previous studies that described roles for these insulin and Tor signaling pathway components in eye development (Chen *et al*. 1996; Bohni *et al*. 1999; Goberdhan *et al*. 1999; Huang *et al*. 1999; Ito and Rubin 1999; Verdu *et al*. 1999; Weinkove *et al*. 1999; Gao *et al*. 2000; Oldham *et al*. 2000; Potter *et al*. 2001).

**Figure 1.**
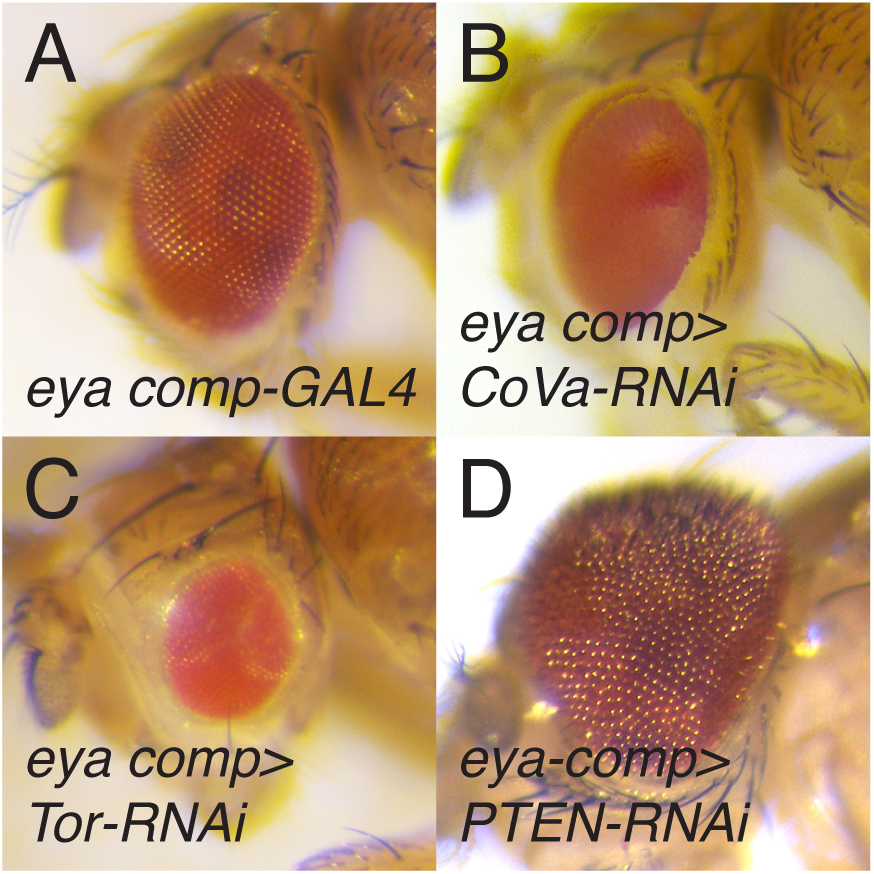
Eye phenotypes caused by RNAi disruption of OXPHOS and growth control. (A) An *eya composite-GAL4/+* control eye (*eya comp-GAL4*). (B) RNAi depletion of *CoVa*, targeted using BDSC 27548, resulted in a glossy-eyed phenotype. (C-D) TRiP RNAi transgenes targeting the growth control regulators (C) Tor, targeted using BDSC 33951, and (D) PTEN, targeted using BDSC 25967, induced small and large eyes, respectively. For all images, *eya composite-GAL4* is abbreviated *eye comp*.

Our ability to identify TRiP lines that interfere with the expression of known growth regulators suggests that our screen efficiently identified key metabolism-associated genes involved in eye development. We would note, however, that a screen of this nature will inevitably produce false-positive results due to the off-target RNAi effect and false negative results due to inefficient depletion of target transcripts. Therefore, we will limit the Results and Discussion sections to those pathways that are either represented by multiple positive results or are notably absent in our analysis.

### Oxidative Phosphorylation

Of the 164 RNAi transgenes that consistently induced an eye phenotype when crossed to the *eya composite*-*GAL4* driver, 40 targeted genes that encode subunits of the ETC and ATP synthase (F-type) as defined by KEGG pathway dme00190 (Figure 2; Supplemental Table 4). These results indicate that eye development is quite sensitive to disruption of Complex I, Complex IV, and Complex V (F-type ATP-synthase), as nearly half of the transgenes that targeted these complexes induced an eye phenotype (Figure 2; Supplemental Tables 2 and 3). In addition, expression of the siRNA that targeted *Coenzyme Q biosynthesis protein 2* (*Coq*2; FBgn0037574; BDSC 53276), which is required for Coenzyme Q production, and *Cytochrome C proximal* (*Cyt-c-p*; FBgn0284248; BDSC 64898) resulted in highly penetrant glossy-eye phenotypes (Figure 2; Supplemental Tables 2-4). We would also note that while expression of only one siRNA associated with Complex II or III induced a phenotype, our screen included relatively few strains that targeted these complexes.

**Figure 2.**
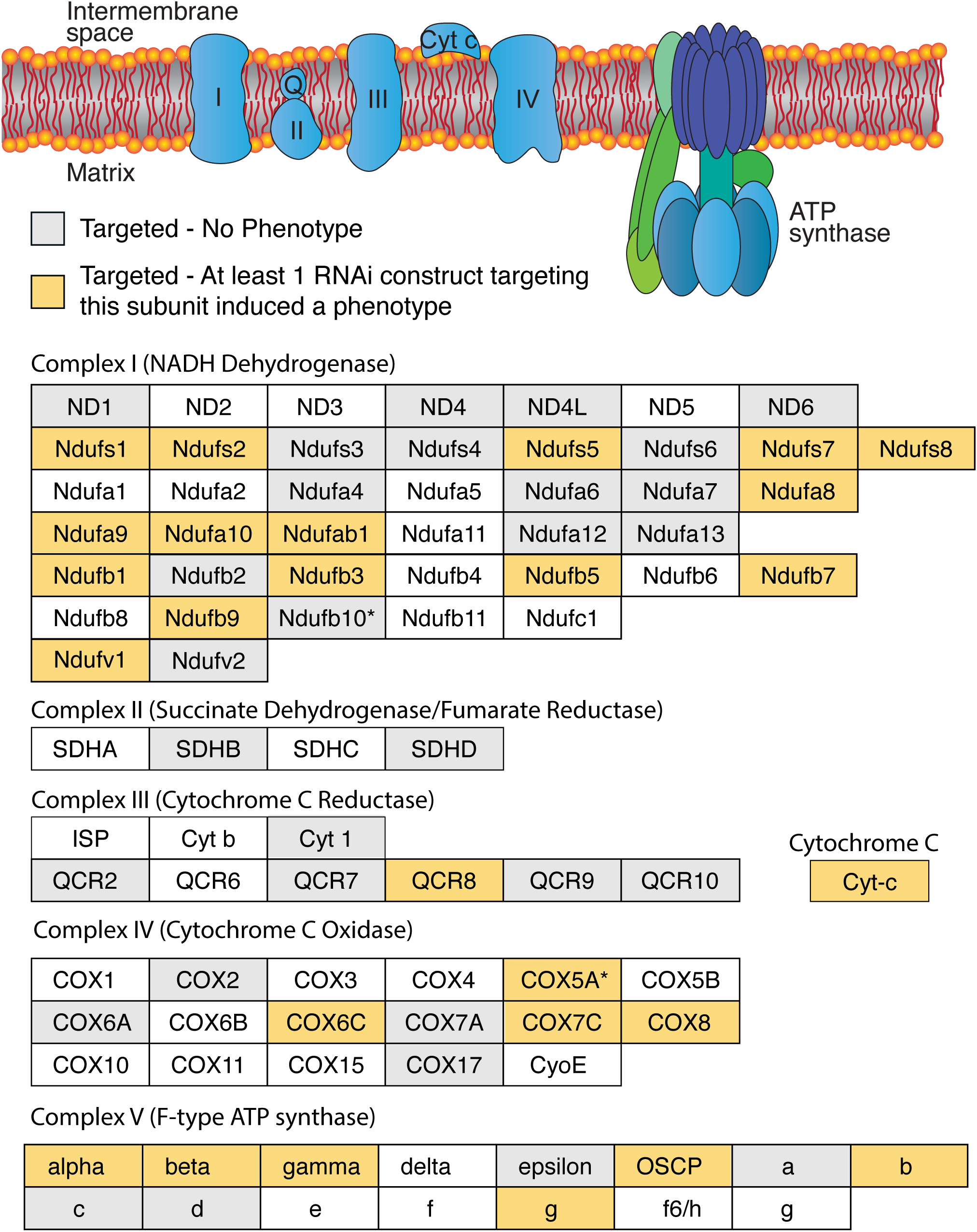
The ETC and ATP synthase are required for normal eye development. (Top) A diagram illustrating the ETC and ATP synthase within the inner mitochondrial membrane. (Below) Individual subunits are listed in boxes and organized by complex. Yellow-shaded boxes indicate that at least one RNAi transgene targeting the subunit induced a phenotype. Grey-shaded boxes indicate that none of the RNAi transgenes targeting this subunit induced a phenotype. Corresponding data can be found in Supplemental Table 4. Diagram is a modified from the illustration presented on the KEGG website for pathway dme00190.

Our findings regarding the ETC and ATP synthase are notable because, among the metabolism-associated TRiP transgenes capable of inducing an eye phenotype, those that disrupt OXPHOS represent one of the largest and most significantly enriched groups (p < 0.0001, Fisher’s exact test, two-tailed, 94 OXPHOS transgenes tested). Moreover, ETC-related transgenes were uniquely associated with the same morphological phenotype - not only did disruption of OXPHOS almost invariably induce a glossy-eye (Figure 3A-C; Supplemental Table 4), but among the RNAi lines tested, almost all of the transgenes that resulted in a glossy-eye phenotype were associated with OXPHOS (Supplemental Tables 2 and 3). Overall, the phenotypic similarities displayed among the OXPHOS-associated TRiP lines support two previously state hypotheses (see Mandal *et al*. 2005; Liao *et al*. 2006): (1) ETC subunits influence eye development in a similar manner. (2) Considering that the function of many nuclear-encoded mitochondrial proteins remain unknown (Pagliarini *et al*. 2008; Pagliarini and Rutter 2013; Calvo *et al*. 2016), targeted disruption of these uncharacterized genes within the eye imaginal disc could potentially identify novel OXPHOS regulators by simply using the glossy-eye phenotype as a readout.

**Figure 3.**
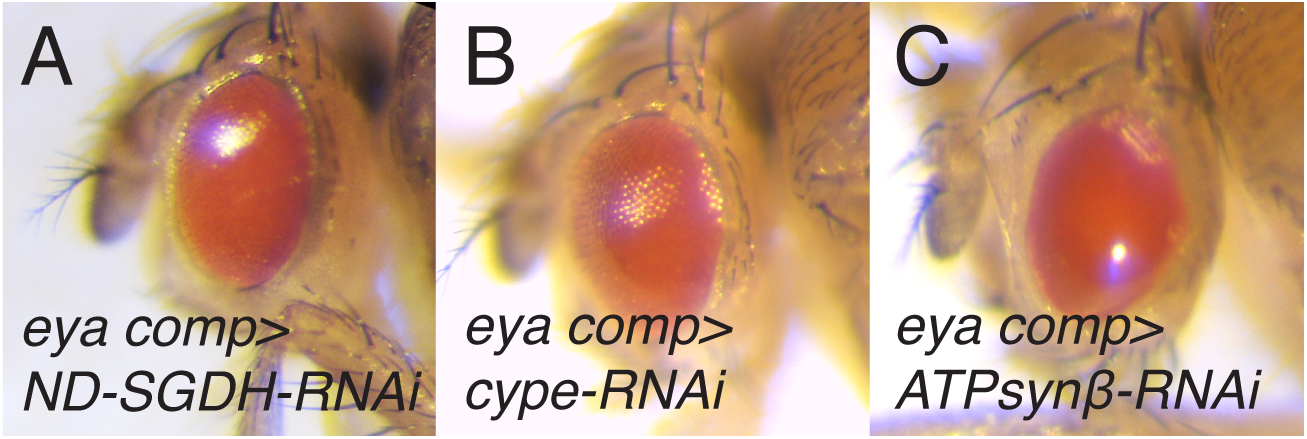
Disruption of the ETC and ATP synthase induces a glossy-eye phenotype. Representative images illustrating how RNAi depletion of OXPHOS components induce a glossy-eyed phenotype. (A) *ND-SGDH*, targeted using BDSC 67311. (B) *cype*, targeted using BDSC 33878. (C) *ATPsyn*b, targeted using BDSC 27712. For all images, *eya composite-GAL4* is abbreviated *eye comp*.

### Glycosylphosphatidylinositol (GPI)-Anchor Synthesis

Many of the enzymes involved in GPI-anchor synthesis emerged as being essential for normal eye development. Of the 8 genes that are associated with this metabolic pathway (KEGG dme00563) and were examined during our screen, TRiP lines that targeted five of these genes induced rough eye phenotypes (Figure 4A-D; Supplemental Table 5). These enzymes represent multiple steps within GPI-anchor biosynthesis, which is consistent with previous observations that this pathway is essential for the function of key proteins involved in eye development, including *rhodopsin, chaoptin*, and *dally* (Krantz and Zipursky 1990; Kumar and Ready 1995; Nakato *et al*. 1995; Satoh *et al*. 2013). Considering that the GPI-anchor biosynthetic enzymes would be predicted to emerge from a screen of this nature, these findings suggest that our approach effectively identified genes involved in eye development.

**Figure 4.**
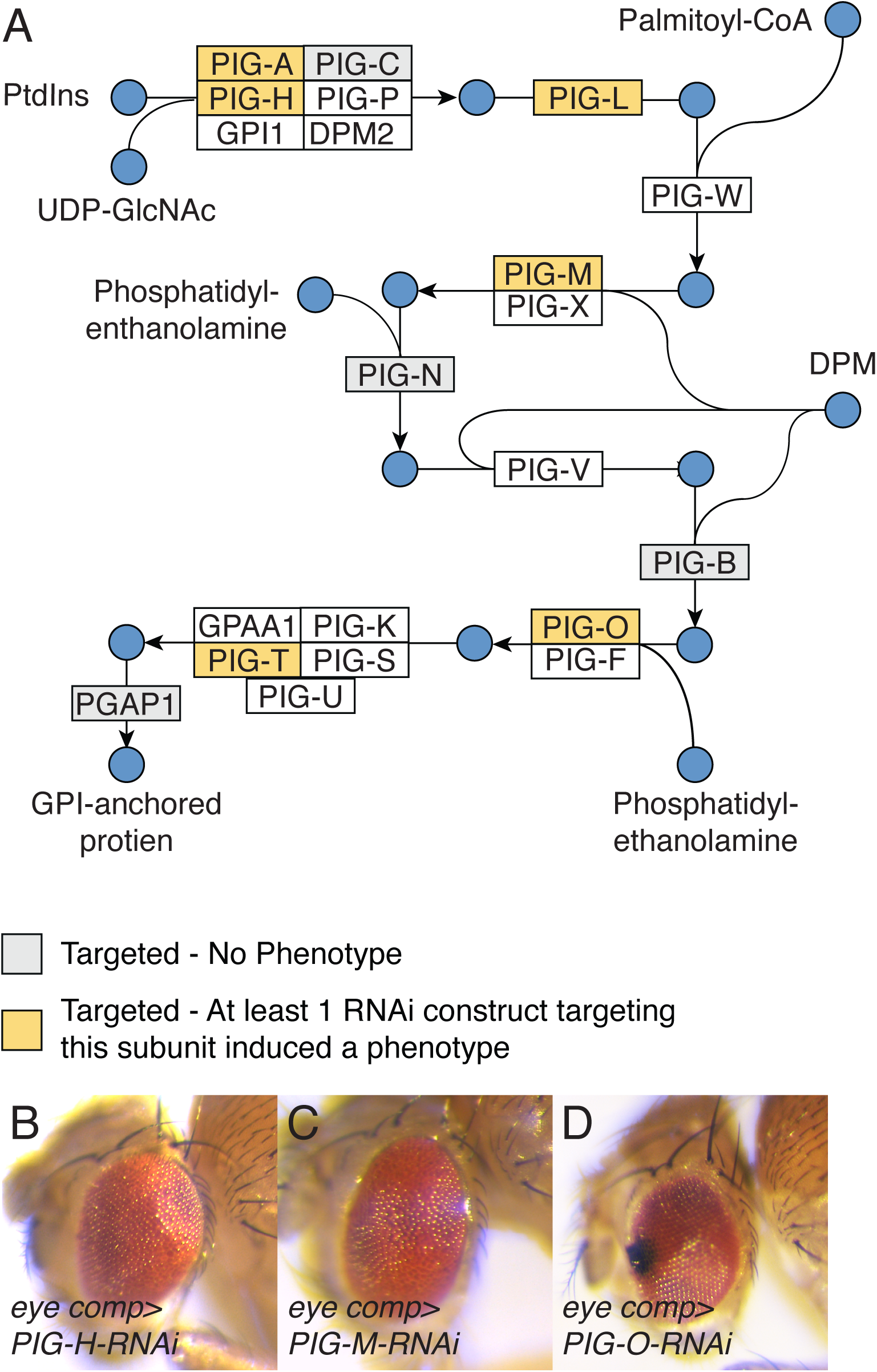
Disruption of GPI-anchor biosynthesis induces a rough eye phenotype. (A) A diagram illustrating GPI-anchor biosynthesis. Diagram is based upon KEGG pathway dme00563. Abbreviations: Phosphatidyl-1D-myo-inositol (PtdIns); Dolichyl phosphate D- mannose (DPM). (B-D) Representative images showing the rough eye phenotype caused by RNAi-induced disruption of (B) *PIG-H*, targeted using BDSC 67330, (C) *PIG-M*, targeted using BDSC 38321, and (D) *PIG-O*, targeted using BDSC 67247. For all images, *eya composite-GAL4* is abbreviated *eye comp*.

### Glycolysis and the TCA cycle

Expression of RNAi constructs targeting either glycolysis (KEGG dme00010) or the TCA cycle (KEGG dme00020) rarely affected eye development. Only two of the 40 TRiP lines that disrupt expression of glycolytic enzymes and none of the transgenes that targeted genes associated with the TCA cycle (n=27) induced an eye phenotype (Supplemental Tables 6 and 7). These results, while surprising, require confirmation using null alleles of these genes, as we can’t eliminate the possibility that enzymes in glycolysis and the TCA cycle are so abundant that RNAi is incapable of reducing their expression below a threshold level. However, we would note that the eyes of *Mitochondrial Pyruvate Carrier 1* (*Mpc1*; FBgn0038662) mutants appear morphologically normal (Supplemental Figure 1B, Bricker *et al*. 2012). Considering that *Mpc1* mutants are unable to transport pyruvate into their mitochondria, eye development must be able proceed normally when glycolysis is uncoupled from the TCA cycle (Bricker *et al*. 2012). Secondly, we previously demonstrated that the TRiP line targeting *Phosphofructokinase* (*Pfk;* FBgn0003071; BDSC 34366) reduces *Pfk* mRNA levels by ∼80%, significantly decreases pyruvate levels, and restricts larval growth (Li *et al*. 2018), however, *Pfk-RNAi* does not interfere with eye development (Supplemental Figure 1C). Although we have not yet confirmed the effectiveness of this *Pfk-RNAi* transgene in the eye imaginal disc, the absence of a phenotype in our screen is notable and warrants future analysis using *Pfk* loss-of-function mutations.

While additional studies are required to understand how glycolysis and the TCA cycle influence eye development, and we doubt that either pathway is completely dispensable in this context, our results raise several intriguing hypotheses. Metabolomic studies of the *Mpc1* mutants reveal that fly larvae raised on standard media readily adapt to this severe disruption of central carbon metabolism (Bricker *et al*. 2012). The same compensatory mechanisms that are activated in *Mpc1* mutants could also support eye development under conditions of reduced glycolytic flux. In addition, considering the apparent dependence of developing eye cells on catabolism of the amino acid glutamine (see below), glucose might not be the primary energy source used by these cells. Finally, we would note that when compared with other larval organs, such as the muscle and brain, imaginal discs express low levels of *Lactate Dehydrogenase* (dLdh, Rechsteiner 1970; Wang *et al*. 2016). Therefore, glycolytic flux appears to be regulated differently in the eye when compared with other larval tissues (Rechsteiner 1970).

### Pentose Phosphate Pathway

Two enzymes within the oxidative shunt of the pentose phosphate pathway (KEGG dme0030), glucose-6-phosphate dehydrogenase (G6PDH; known as *Zwischenferment*; FBgn0004057; BDSC 50667) and phosphogluconate dehydrogenase (*Pgd*; FBgn0004654; BDSC 65078) produced a small eye phenotype (Supplemental Tables 2 and 3). These results were unexpected because both enzymes are thought to be dispensable for growth and viability - *Zw* mutants display no discernable phenotype, and while *Pgd* mutants are lethal, *Zw Pgd* double mutant are viable with no obvious morphological defects (Hughes and Lucchesi 1977). While such results require confirmation using clonal analysis, our observations hint at the possibility that tissue-specific disruption of the pentose phosphate pathway can induce developmental phenotypes – a phenomenon that has been previously observed in studies of *Drosophila* metabolism (Caceres *et al*. 2011). Considering that the oxidative branch of the pentose phosphate pathway serves a key role in maintaining NAPDH levels (Ying 2008), future studies should examine the possibility that eye development relies on G6PDH and PGD to maintain this pool of reducing equivalents.

### Glutamine metabolism

Our screen revealed an unexpected role for glutamine (Gln) and glutamate (Glu) in eye development. Of the 24 TRiP lines that targeted genes directly involved in Gln/Glu metabolism (see enzymes that interact with Gln/Glu in KEGG pathway dme00250), five induced either a small or no eye phenotype (Supplemental Table 8). These five RNAi lines targeted five genes that directly regulate Gln/Glu-dependent metabolic processes (Figure 5A):

(1) *bb8* (FBgn0039071; BDSC 57484) encodes the enzyme glutamate dehydrogenase (GLUD), which is responsible for converting glutamate into a-ketoglutarate and ammonia (see KEGG dme00250). Because GLUD can funnel glutamate into the TCA cycle, this enzyme allows cells to generate both fatty acids and ATP in a glucose-independent manner – as is evident by the fact that many cancer cells adapt to inhibition of glycolysis by up-regulating GLUD activity (Altman *et al*. 2016). A recent study in *Drosophila* has implicated *bb8* in promoting spermatogenesis (Vedelek *et al*. 2016).
(2) *CG8132* (FBgn0037687; BDSC 57794) encodes an omega-amidase that is homologous to the human nitrilase family member 2 (NIT2) enzyme, which converts a-ketoglutaramate into a-ketoglutarate and ammonia (Jaisson *et al*. 2009; Krasnikov *et al*. 2009). The endogenous function of this enzyme remains poorly understood in animal systems, however, there are some indications that NIT2 functions as a tumor suppressor in humans (Zheng *et al*. 2015).
(3) Glutamine synthetase 1 (Gs1; FBgn0001142; BDSC 40836) generates Gln from ammonia and Glu (Caizzi and Ritossa 1983). Since Gln is required for several biosynthetic processes, including the production of nucleotides, glutathione, and glucosamine-6- phosphate (see below, Altman *et al*. 2016), Gs1 ensures that growing and proliferating cells have adequate levels of this amino acid. In *Drosophila*, this enzyme is also required for early mitotic cycles within syncytial embryos (Frenz and Glover 1996).
(4) Glutamine:fructose-6-phosphate aminotransferase 2 (Gfat2; FBgn0039580; BDSC 34740) converts Gln and fructose-6-phosphate into Glu and glucosamine-6-phosphate (Graack *et al*. 2001). In turn, glucosamine-6-phosphate is used to generate *N-*acetyl-glucosamine (GlcNAc), which is required for several cellular processes, including chitin formation and protein modifications. Moreover, the multifaceted roles for glucosamine-6-phosphate and GlcNAc in development are essential for cell proliferation and tissue growth, as demonstrated by the recent observation that *Drosophila Gfat2* is required for proliferation of adult intestinal stem cells (Mattila *et al*. 2018).
(5) *phosphoribosylamidotransferase* (*Prat*; FBgn0004901l; BDSC 43296) links Gln with purine biosynthesis (Clark 1994). RNAi targeting of this gene in the eye imaginal disc resulted in a lethal phenotype during our initial screen (Supplemental Table 2), suggesting that disruption of nucleotide production in the developing eye induces non-autonomous effects. Our observation is consistent with previous results, which indicate that *Prat* is expressed in L3 imaginal discs and that *Prat-RNAi* results in a pupal lethality (Ji and Clark 2006; Brown *et al*. 2014).

**Figure 5.**
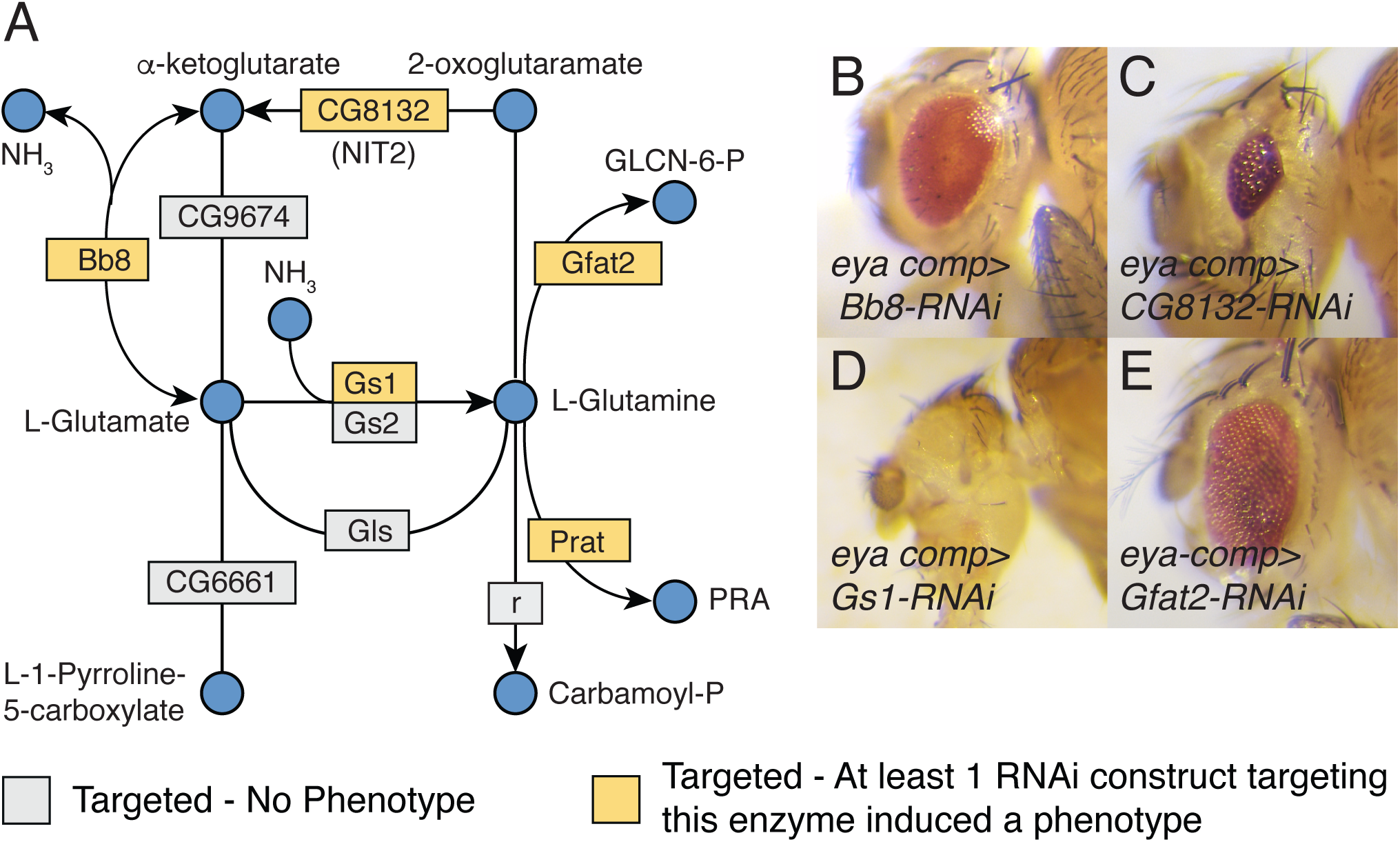
Enzymes associated with glutamate (Glu) and glutamine (Gln) metabolism are essential for normal eye development. (A) A diagram illustrating the metabolic reactions associated with Glu and Gln metabolism as defined by KEGG pathway dme00250. (B-E) Representative images illustrating how disruption of Glu/Gln metabolism affects eye development. Abbreviations: D-glucosamine-6-phosphate (GLCN-6-P) and 5- phosphoribosylamine (PRA). (B) *Bb8*, targeted using BDSC 57484. (C) *CG8132*, targeted using BDSC 38321. (D) *Gs1*, targeted using BDSC 40836. (E) *Gfat2*, targeted using BDSC 34740. For all images, *eya composite-GAL4* is abbreviated *eye comp*.

In addition to the enzymes that are directly involved in Glu/Gln catabolism, RNAi of two additional genes associated with these amino acids elicited eye phenotypes:

(1) γ-glutamyl transpeptidase (*Ggt-1*; FBgn0030932; BDSC 64529) transfers a γ-glutamyl residue from a donor molecule, such as glutathione, to an acceptor molecule (Ikeda and Taniguchi 2005; Heisterkamp *et al*. 2008). Moreover, this enzyme can generate Glu by using water as an acceptor molecule for γ-glutamyl (Ikeda and Taniguchi 2005; Heisterkamp *et al*. 2008). *Drosophila Ggt-1* was previously reported to function in the larval Malpighian tubules, where it facilitates green-light avoidance by generating glutamate (Liu *et al*. 2014).
(2) *Selenide water dikinase* (*SelD*; FBgn0261270; BDSC 29553) encodes a member of the selenophosphate synthetase 1 (SPS1) enzyme family that does not synthesize selenophosphate but rather functions in redox homeostasis and glutathione metabolism (Xu *et al*. 2007a; Xu *et al*. 2007b; Tobe *et al*. 2016). Consistent with the proposed functions of SPS1 proteins, *SelD* serves a critical role in *Drosophila* eye development by restricting reactive oxygen species (ROS) accumulation (Morey *et al*. 2003). In the absence of SelD function, elevated ROS levels within the eye interfere with a variety of developmental signaling events (Alsina *et al*. 1999; Morey *et al*. 2001), as evident by the fact that the *SelD* null mutation *patufet* (*SelD^ptuf^*) dominantly suppresses the eye and wing phenotypes induced by ectopic activation of *sevenless* and *Raf* (Morey *et al*. 2001). While the exact metabolic function of SelD remains unknown, *SelD* knockdown in SL2 cells induces excessive Gln accumulation (Shim *et al*. 2009).

We find these results notable because these seven enzymes are involved in a diversity of metabolic processes, including biosynthesis, energy production, and cell signaling. Moreover, not only are many of these enzymes implicated in cancer cell proliferation and tumor growth (Lin *et al*. 2007; Altman *et al*. 2016), but one of the metabolites associated with these enzymes, a-ketoglutarate, is an essential regulator of histone methylation and gene expression (Chisolm and Weinmann 2018). Overall, our findings indicate that *Drosophila* eye development could serve as a powerful *in vivo* model for investigating how Glu/Gln metabolism influences cell proliferation and tissue growth.

## DISCUSSION

Here we use the *Drosophila* TRiP RNAi collection to identify metabolic processes that are required for the growth and development of the eye. Our screen not only verified that RNAi could effectively disrupt metabolic processes with known roles in eye development (e.g., *CoVa*, ETC subunits, enzymes involved in GPI-anchor biosynthesis), but also proved effective at identifying additional pathways that are essential for the growth of this tissue. Here we highlight two key findings that we believe warrant further examination.

### Metabolic pathways are associated with specific developmental events

The RNAi phenotypes uncovered in our screen demonstrate how different stages in eye development impart unique demands on intermediary metabolism. For example (and as previously described by the Banerjee lab), the OXPHOS-associated glossy eye phenotype results from a cell cycle arrest during the second mitotic wave (Mandal *et al*. 2005), resulting in the loss of pigment cells and lens secreting cone cells (for a review of cone and pigment cell development, see Kumar 2012). A key feature of this phenotype is that the overall eye size remains normal, indicating that OXPHOS disruption does not interfere with cell proliferation ahead of the morphogenetic furrow. The unique nature of this phenotype suggests that any TRiP line inducing a glossy, normal sized eye should be investigated for a potential role in OXPHOS. Similarly, the rough eye phenotype induced by RNAi of GPI-anchor biosynthesis likely reflects the disruption of proteins required for the formation of ommatidium, including those associated with morphogen signaling, cell polarity, and cell specification (Kumar 2012). Therefore, those genes associated with a rough eye phenotype in our screen should be examined for potential roles in ommatidial assembly.

While our screen indicates that dozens of metabolic enzymes are required for eye development, perhaps our most intriguing results are the small/no eye phenotypes induced by the disruption of Glu/Gln metabolism. These developmental defects likely stem from either decreased cell proliferation ahead of the morphogenetic furrow or defects in cell fate specification (for review, see Kumar 2011) and are consistent with the role of Glu/Gln-associated enzymes in mammalian cell proliferation and differentiation (Altman *et al*. 2016). The developing eye disc, therefore, provides an ideal model to understand how signal transduction cascades regulates Glu/Gln metabolism and investigate how the metabolism of these amino acids influence cell proliferation and tissue growth. Moreover, considering that relatively few TRiP lines induced either a small eye or no eye, any gene associated with this phenotype should be examined for links with Glu/Gln metabolism. For example, RNAi targeting the *Drosophila* gene *jet fuel* (FBgn0033958; BDSC 43284) induces a small eye phenotype. This gene, which encodes a major facilitator superfamily transporter protein involved in nociception (Honjo *et al*. 2016), is uncharacterized during eye development. Based on the phenotypes observed in our screen, future studies should examine potential links between *jet fuel* function and Glu/Gln metabolism.

### The *Drosophila* eye as a model for studying metabolic plasticity and robustness

Our screen further supports previous observations that *Drosophila* development is surprisingly resistant to metabolic insults. Our observation that eye development was largely unaffected by the RNAi transgenes that target glycolysis and the TCA cycle was unexpected. While we doubt that either pathway is completely dispensable for eye formation, our results are consistent with the ability of *Drosophila* development to withstand severe metabolic insults (e.g., *Mpc1* mutants, Bricker *et al*. 2012). This metabolic robustness makes sense because animal development must readily adapt to a variant of nutrient sources and environmental stresses. Based upon the results of this screen, we propose that the fly eye could serve as a model to identify the compensatory pathways that that allow cell growth and proliferation to proceed in the face of major metabolic disruptions.

Overall, our genetic screen demonstrates how *Drosophila melanogaster* can serve as a powerful model to identify tissue-specific metabolic factors required for tissue growth and organogenesis. Moreover, we believe this work represents a necessary step toward systematically analyzing the metabolic pathways that support cell proliferation and tissue growth within the fly.

## Supporting information

Suppemental Table 3

Suppemental Table 4

Suppemental Table 5

Suppemental Table 6

Suppemental Table 7

Suppemental Table 8

## ACKNOWLEDGEMENTS

We thank the TRiP at Harvard Medical School (NIH/NIGMS R01-GM084947) for providing transgenic RNAi fly stocks used in this study. We also thank the Bloomington *Drosophila* Stock Center (NIH P40OD018537), Cale Whitworth for helping us search the BDSC database, Flybase (NIH 5U41HG000739), and the members of the Kumar lab for providing reagents and technical support. Thanks to Kudakawashe Tshililiwa, Curteisha Jacobs, and Joy Morounfolu for assistance with genetic crosses. Special thanks to Bonnie Weasner and Justin Kumar for support, advice, helpful discussions, and a critical reading of the mansucript. J.M.T. is supported by the National Institute of General Medical Sciences of the National Institutes of Health under a R35 Maximizing Investigators’ Research Award (MIRA; 1R35GM119557).

---

**Supplemental Figure 1.**
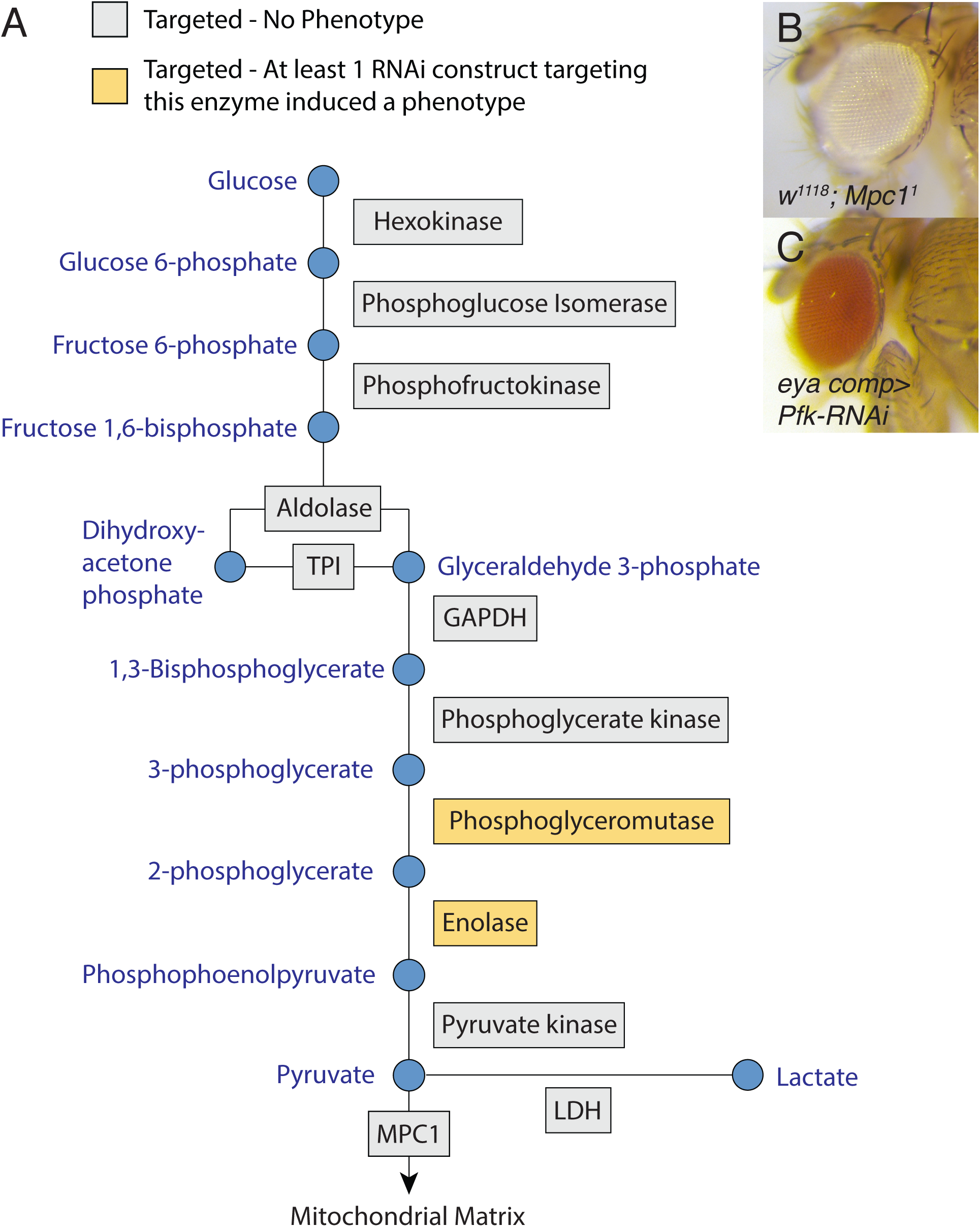
Most TRiP RNAi lines that target glycolysis do not disrupt eye development. (A) All enzymatic steps in glycolysis were targeted during the course of the screen. Yellow-shaded boxes indicate that at least one RNAi transgene targeting the enzyme induced a phenotype. Grey-shaded boxes indicate that none of the RNAi transgenes targeting this subunit induced a phenotype. Corresponding data can be found in Supplemental Table 6. Diagram is modified from the pathway illustrating KEGG pathway dme00010. (B) *w^1118^; Mpc1^1^* mutant eyes are morphologically normal, indicating that glucose oxidations is not required during eye development. (C) RNAi targeting *Pfk* using BDSC 34366 failed to induce an eye phenotype. *eya composite-GAL4* is abbreviated *eye comp*.

**Supplemental Figure 2.**
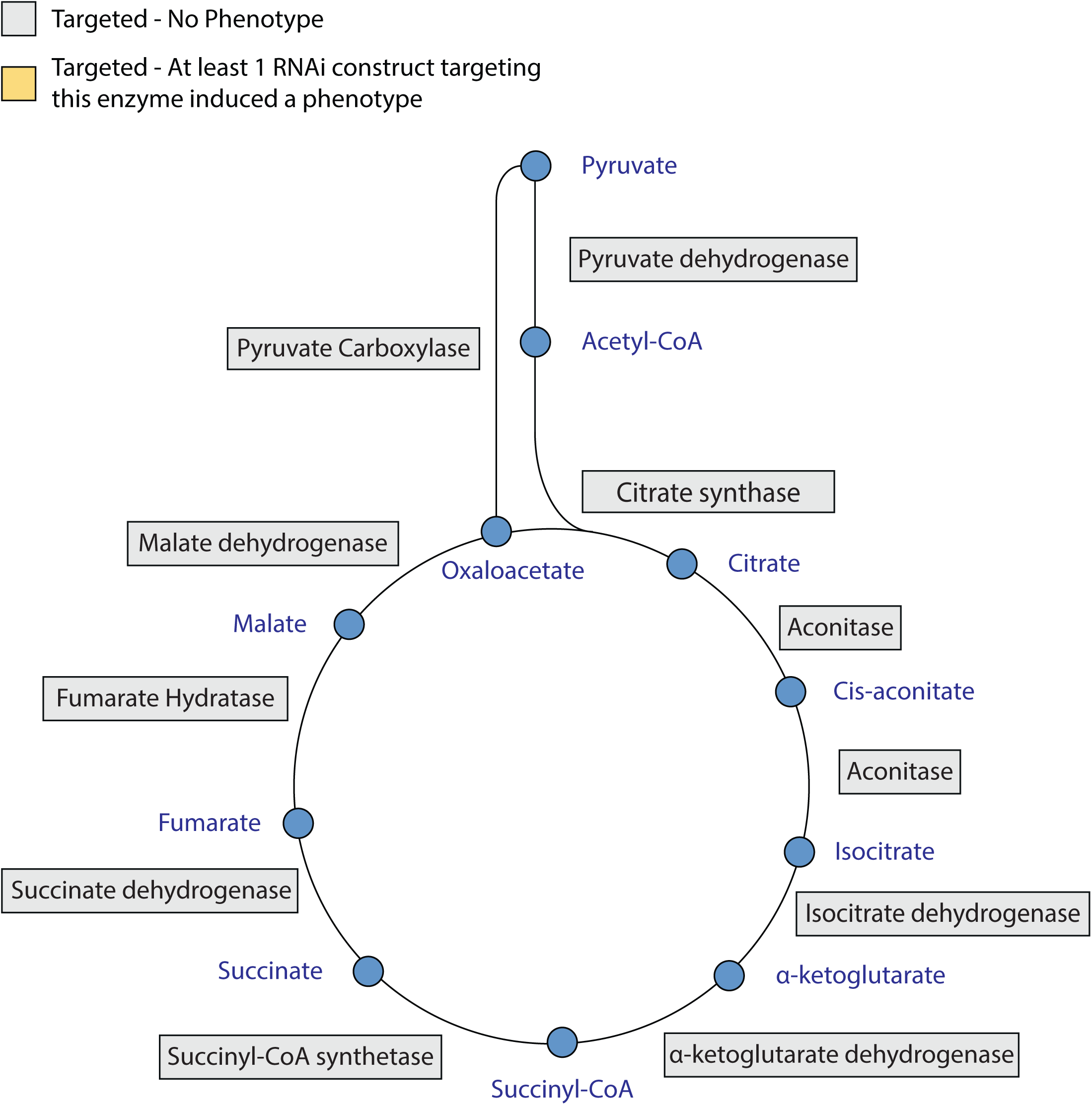
TRiP RNAi lines targeting the TCA cycle do not disrupt eye development. Grey-shaded boxes indicate that none of the RNAi transgenes targeting this enzyme induced a phenotype. Corresponding data can be found in Supplemental Table 7. Diagram is a modified from the pathway illustrating KEGG pathway dme00020.

**Supplemental Table 1.** A list of *Drosophila* genes involved in metabolism and nutrient sensing

**Supplemental Table 2.** Results of TRiP RNAi metabolism screen_Trial 1

**Supplemental Table 3.** Results of TRiP RNAi metabolism screen_Trial 2

**Supplemental Table 4.** A list of the BDSC TRiP RNAi strains that were used to disrupt oxidative phosphorylation

**Supplemental Table 5.** A list of the BDSC TRiP strains that were used to disrupt GPI-anchor biosynthesis.

**Supplemental Table 6.** A list of the BDSC TRiP strains that were used to disrupt glycolysis.

**Supplemental Table 7.** A list of the BDSC TRiP strains that were used to disrupt the TCA cycle.

**Supplemental Table 8.** A list of the BDSC TRiP strains that were used to disrupt glutamate and glutamine metabolism.

